# A Biologically Plausible Model for Continual Learning using Synaptic Weight Attractors

**DOI:** 10.1101/2021.10.20.465035

**Authors:** Romik Ghosh, Dana Mastrovito, Stefan Mihalas

## Abstract

The human brain readily learns tasks in sequence without forgetting previous ones. Artificial neural networks (ANNs), on the other hand, need to be modified to achieve similar performance. While effective, many algorithms that accomplish this are based on weight importance methods that do not correspond to biological mechanisms. Here we introduce a simple, biologically plausible, method for enabling effective continual learning in ANNs. We show that it is possible to learn a weight-dependent plasticity function that prevents catastrophic forgetting over multiple tasks. We highlight the effectiveness of our method by evaluating it on a set of MNIST classification tasks. We further find that the use of our method promotes synaptic multi-modality, similar to that seen in biology.

## 1 Introduction

Deep learning models such as artificial neural networks have become a staple for researchers and engineers alike – facilitating advances in a variety of supervised learning tasks. The most impressive models in this area tend to follow the traditional supervised learning format. Specifically, given a dataset *D*_*tr*_ = {(*x*_*i*_, *y*_*i*_)|*i* ∈ *N*} such that *x*_*i*_ denotes a feature vector sampled from the space *X* and *y*_*i*_ denotes a corresponding target vector from the space *Y*, learn a mapping *f*_*θ*_ : *X* ↦ *Y* that minimizes a loss, e.g. 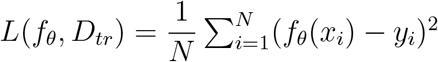 such that *θ* is the set of parameters for a neural network. Deep learning models parameterizing *f* have been incredibly successful in domains ranging from facial recognition to automatic handwriting identification (Taigman et al., 2014; Graves and Schmidhuber, 2009).

An important question to ask is whether this problem formulation readily lends itself to learning multiple tasks in sequence. That is, given a set of tasks τ and datasets *D*_𝒯_, is minimizing the loss 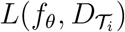 for each task in sequence the same as minimizing the joint loss *L*(*f*_*θ*_, *D*_𝒯_)? Following the work of Goodfellow et al. (2015) it became clear this is not the case. Indeed, modern artificial neural networks struggle to learn multiple tasks in sequence without forgetting previous knowledge, a phenomenon termed *catastrophic forgetting* (Goodfellow et al., 2015).

How to solve this problem has been the subject of an emerging field, of continual learning. We will only describe in the introduction the set of methods related to our approach, but for a more in-depth description of other methods, including regularization and network expansion methods, we recommend a recent review (Parisi et al., 2019).

More complex synaptic plasticity models which include a additional variables associated with a synapse have been proposed (Benna and Fusi, 2016; Kirkpatrick et al., 2017; Zenke et al., 2017). Among them, weight importance methods are particularly adept at increasing retention across a large number of tasks (Kirkpatrick et al., 2017; Zenke et al., 2017). As the name suggests, weight importance methods aim to penalize changes to the weights that contribute most to task accuracy on previous tasks – thereby preventing weight changes that would result in the loss of previously learned information. To date, the most successful weight importance methods are Synaptic Intelligence (SI) and Elastic Weight Consolidation (EWC). EWC imposes a quadratic penalty on the distance from the previous weights proportional to the corresponding location on the diagonal of the Fisher information matrix near the previous minimum. In doing so, EWC attempts to estimate the importance of each parameter as inversely proportional to the Laplace approximation of its expected posterior variance. While effective, the method is expensive and requires frequent recomputation of this diagonal, a process whose cost is directly proportional to the number of outputs for a given task (Kirkpatrick et al., 2017). Alternatively, SI aims to compute a per-parameter regularization strength based on a discrete approximation of the parameter’s previous contribution to decreases in loss (Zenke et al., 2017).

Despite their success, neither SI nor EWC represent compelling biological mechanisms for continual learning. Both the EWC and SI papers correctly make the argument that there is biological evidence for synaptic protection (Yang et al., 2009). However, their methods for computing weight importance either (1) require the computation of a complicated information matrix near a previous task minimum (as in EWC) or (2) require the addition of extra synaptic dimensions in which parameters’ importance is stored (as in SI) (Kirkpatrick et al., 2017; Zenke et al., 2017).

Unlike their artificial counterparts, humans are excellent at learning tasks in sequence. We routinely learn multiple languages and our proficiency in one does not imply that we have forgotten the others. It is well known that humans have evolved both genomic and connectomic priors that can intelligently guide learning dynamics (Zador, 2019). We propose a simple prior a fixed function that maps synapse strength to learning rate. In contrast to SI and EWC, we show that it is not necessary to compute weight importance explicitly to alleviate catastrophic forgetting. Instead, we find that it is possible to meta-learn parameters for this weight-dependent learning rate function. We further show that this prior mapping can effectively prevent catastrophic forgetting in ANNs on sequences of supervised learning tasks. Finally, we find that this method allows the network to approximate the synaptic consolidation performed by SI and EWC (Kirkpatrick et al., 2017; Zenke et al., 2017).

## 2. Methods

We wanted to develop a continual learning method that is both simple and reasonably effective. Moreover, we hoped to develop an algorithm for which there are already potential biological mechanisms. A weight-dependent plasticity function fit all of these requirements. Neuronal plasticity is known to be activity dependent (Cingolani et al., 2008). As such, a weight-dependent plasticity function is a simple and biologically plausible analog. Specifically, we propose a set of algorithms in which we optimize hyperparameters *ϕ* ={a_1_, µ, σ, γ, ζ, λ} over an arbitrary number of tasks for a learning rate function f (W) which takes weights as input and outputs a learning rate multiplier.

To design the function *f*, we draw inspiration from biology. Past work suggests that long-term memories are stored in stable synaptic networks (Yang et al., 2009). To induce this stability, the learning rate function needs to approach zero or dip for some values of the network parameters *θ*. In this way, the function can trap weights in an important region; this is illustrated in *Figure 1b*. It is important that the base learning rate be parameterized as well, such that the network is allowed to learn more quickly in certain weight ranges. To these ends, we chose *f* to be a negative Gaussian function with an added multiplier/offset parameter. Specifically,

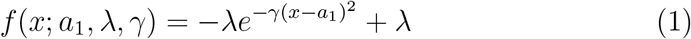

such that *a*_1_, *λ*, and *γ* are hyperparameters describing the location, depth, and width of the dip – respectively. To allow weights to fall into these dips, we only apply *f* to parameters with gradients going away from *a*_1_. Crucially, if *λ >* 1, weights with gradients leading away from the dip may be pushed away more quickly than they otherwise would have been. This allows for the creation of a second group of strongly inhibitory weights that can help sparsify the heavy activity induced by the pooling of positive weights at *a*_1_.

**Figure 1:**
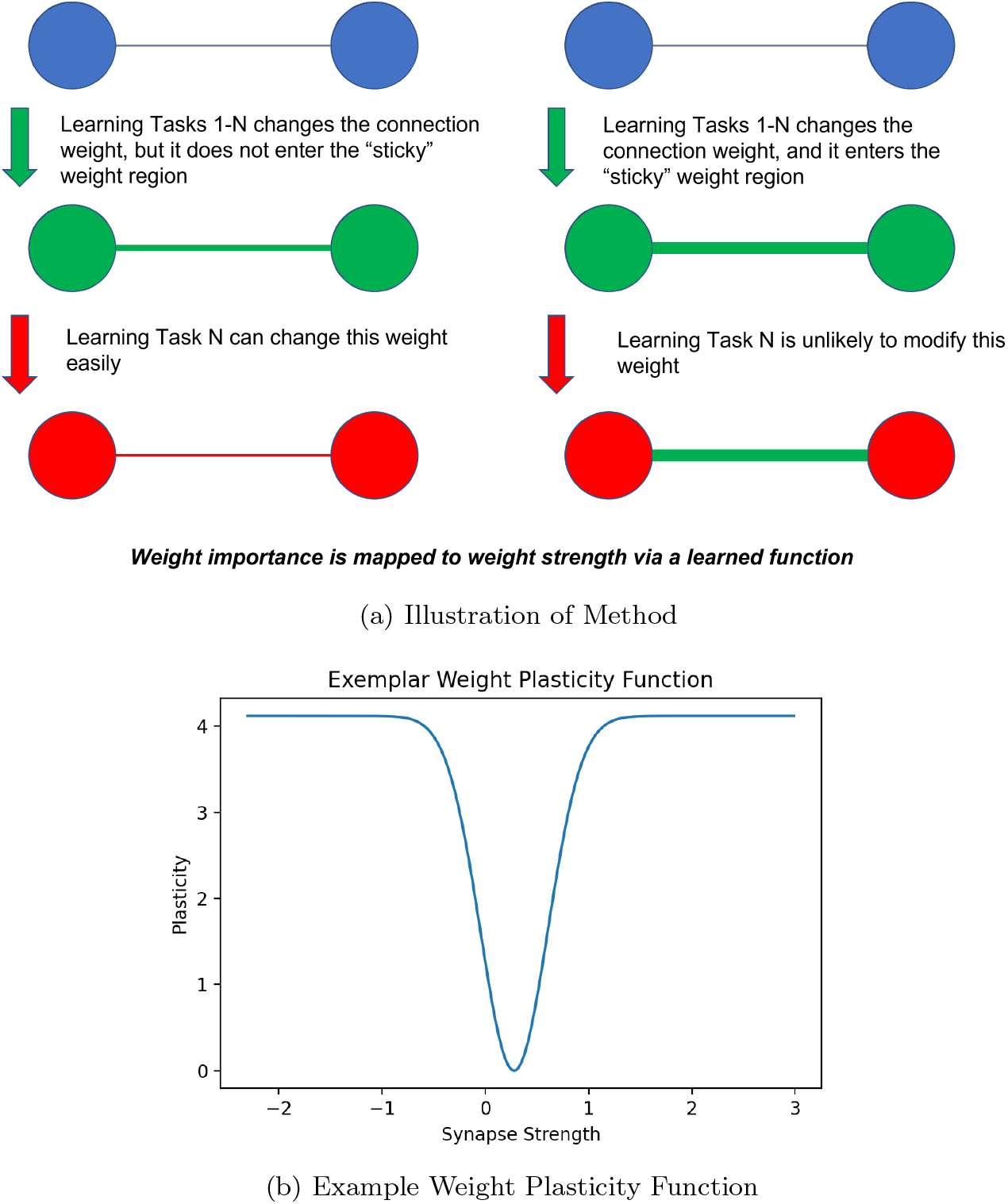
(a) Graphical illustration of the effect of our algorithm; our method traps weights close to attractor values and allows movement of weights that are far away (b) Example plasticity function – remains constant at around 4 and dips to zero near 0.279

It is important that this type of regularization not override the ability of the network to learn. In essence, a good learning rate function *f* traps as few parameters as is necessary to remember the previous task but leaves the rest free to learn representations for new tasks. To affect this dynamic, we add L1 regularization to the system – decreasing gradients towards the well. We provide a visualization of the function in *Figure 1b*.

Finally, we propose a modified update rule. We compute the learning rate multiplier dictated by the function *f* and parameterized by *ϕ* and apply it to the classification gradients going away from *a*_1_.(*Algorithm 1*)

### Algorithm 1

Sticky Gradient

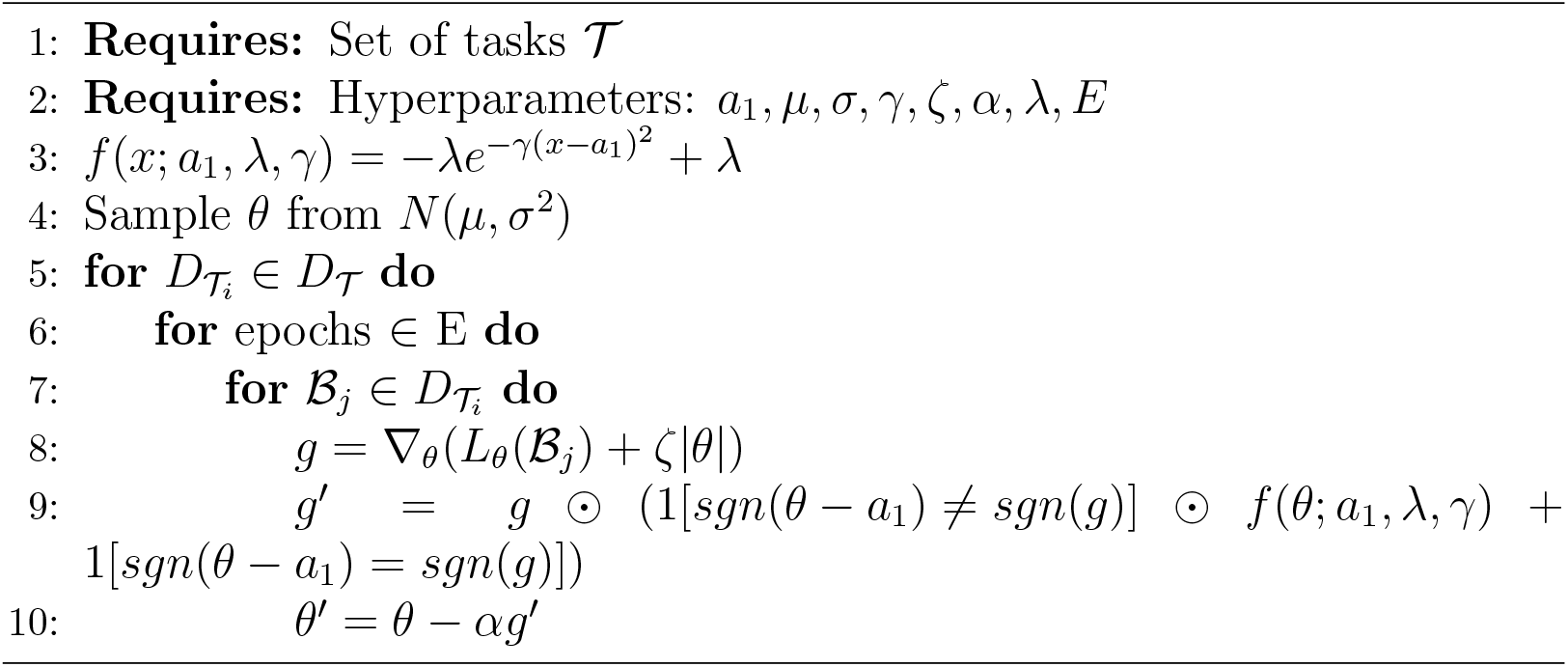

### 2.1. Meta-learning

To learn the parameters *ϕ*, we employ a simple Tree-Structured Parzen Estimator (TPE) to intelligently search over parameter space (Bergstra et al., 2011). To prioritize many-task retention, we use a modified meta-objective function in which averages from later tasks are weighted more than those of earlier tasks. Suppose that *θ*_*j*_ are the parameters of a neural network after being trained on task *j*. Suppose further that 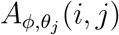 denotes validation accuracy on the *i*^*th*^ task after being trained on the *j*^*th*^ task with learnable hyperparameters *ϕ* and network state *θ*_*j*_. Let *N* be the total number of tasks.

Then the optimization problem can be formulated as

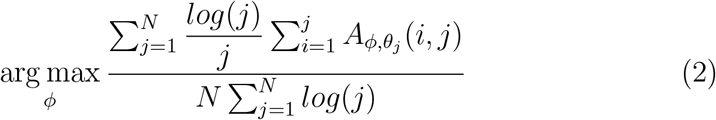

We optimize the model on an arbitrary set of ten classification tasks, described in the following section. Remarkably, we find that it takes as few as 30 trials to find an effective set of parameters, given by *Table 1*.

**Table 1:** Learned hyperparameters from 30 trial meta-search. Surprisingly, the metaoptimization yields a large value of *λ*, implying different learning rate for gradients going away from the attractor is important. The function *f* with these parameters is visualized in *Figure 1b*.

| $a_1$ | $\mu$ | $\sigma$ | $\gamma$ | $\zeta$ | $\lambda$ |
| --- | --- | --- | --- | --- | --- |
| 0.279 | 0.00363 | 0.0204 | 4.70 | 0.000574 | 4.12 |

## 3. Experiments

### 3.1. Permuted MNIST

We use permuted MNIST (PMNIST), a simple variant of the MNIST dataset to evaluate our method in a multi-task setting. PMNIST is simply a derivative of the MNIST dataset in which the handwritten images are normalized, flattened, and randomly permuted n-times such that n is the desired number of tasks (Goodfellow et al., 2015).

### 3.2. EWC Comparison

To establish the value of this method, we compare it to EWC on the permuted MNIST benchmark (Kirkpatrick et al., 2017). We use a simple feed-forward architecture with a single ReLU hidden layer consisting of 2000 units and instantiate three networks to be trained using EWC with a penalty multiplier of 1000, the *sticky gradient* method, or a vanilla learning algorithm (control) (Kirkpatrick et al., 2017). All networks are trained for a single epoch on ten tasks, different than those used in the meta-optimization process, and optimized using Adam with a base learning rate of 1e-4. It is important to note that we sample a new set of ten tasks for each random seed utilized but do not alter *ϕ*. That is, the meta-parameters remain constant across different samples of 10 tasks from PMNIST.

Impressively, our method outperforms the control by an average margin of around 20% after 10 tasks, indicating a significant advantage in its retentive ability (*Fig. 2a*). While EWC outperforms our method by an average accuracy of 2.5% at ten tasks, we observe that this difference decreases across tasks (*Fig. 2a*). Our method underperforms EWC on a test set containing all previous tasks by about 2.7%. However, our method retains the some ability to learn new tasks, enjoying a 3% benefit on the most recent task at 10 tasks (*Fig. 2c*). Of course, the control maintains nearly all of its flexibility at the cost of an almost complete lack of retention, 8% better than our method and 11% better than EWC on the ten task horizon.

**Figure 2:**
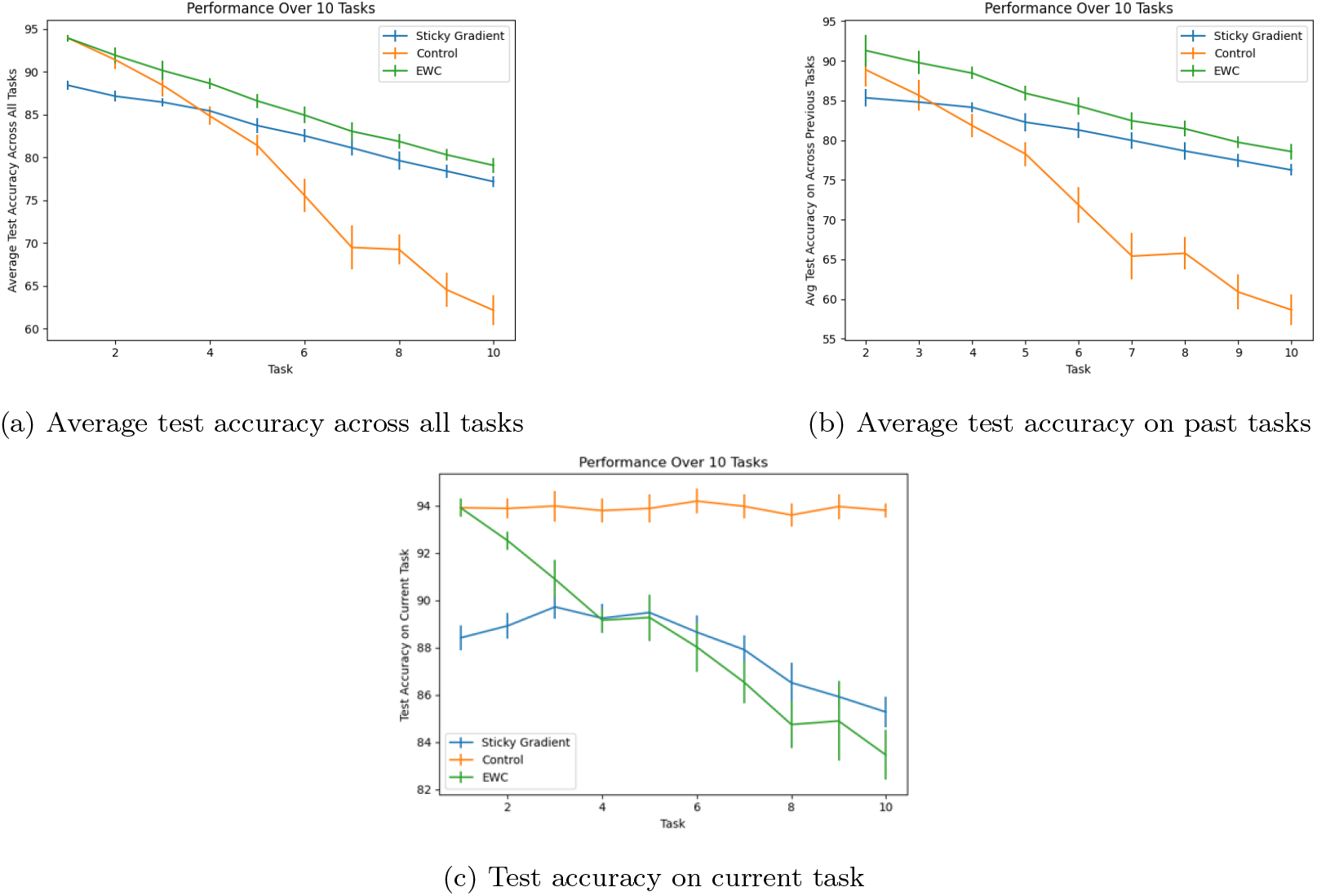
(a) Comparison study detailing the average test accuracy of trials under 10 different random seeds. Accuracy on all tasks is recomputed after every task on a held-out test set and averaged; error bars denote 95% confidence intervals. EWC and the *sticky gradient* method outperform the control by an average margin of around 19%, but EWC outperforms our method by an average margin of 2.5 % at ten tasks. (b) Average test accuracy on all tasks, not including the current one. Accuracy is computed on all previous tasks after every task except the first and averaged; error bars are 95% confidence intervals. Our method lags further behind EWC in this metric – at around a 2.7% performance penalty, but significantly outperforms the control. (c) Test accuracy on current task; error bars denote a 95% confidence interval. Our method and EWC have a decreased ability to learn new tasks; both lag the control’s current task performance by around 9% at ten tasks, as the networks become restricted in learning by maintaining the previous tasks.

### 3.3. Ablation Studies

Given that our method makes use of an L1 regularization and smart initialization – we find it necessary to disentangle the effects of each of these sub-procedures from the whole. Controlling for *ϕ*, we test every combination of L1 regularization, the *sticky gradient* method, and initialization distribution. We denote network initialization from a learned normal distribution, parameterized by *µ* and *σ*, as *smart initialization*. All networks are simple feed-forward MLPs with a single hidden layer of 2000 units coupled with ReLU activations, trained for a single epoch, and optimized by Adam with a base learning rate of 1e-4.

We find that the *sticky gradient* method, coupled with both the L1 regularization and *smart initialization* has the best average accuracy at 10 tasks around 3% better than its closest competitor, a combination of the *sticky gradient* and *smart initialization*, and about 6% better than its second closest competitor – simple *smart initialization*. Methods with the *sticky gradient* and *smart initialization* retain a substantial benefit over the control throughout the training process, starting at around four tasks (*Fig. 3*). For brevity, we now refer to the combination of the *sticky gradient*, L1 regularization, and smart initialization as the *sticky gradient* method.

**Figure 3:**
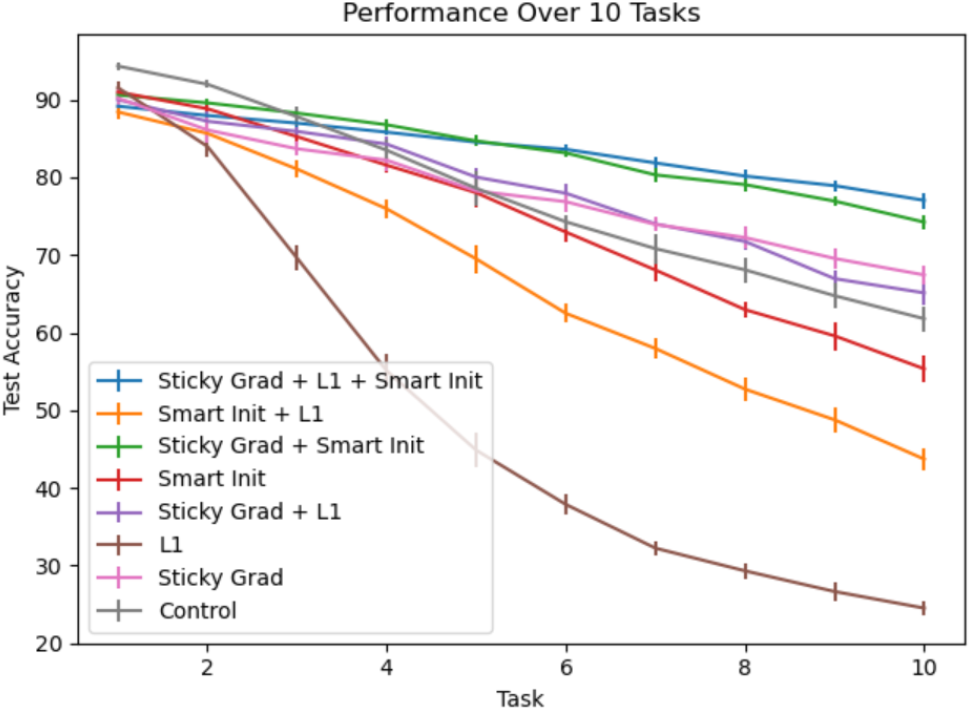
Ablation Study detailing the test accuracy of trials of trials under 10 different random seeds. Accuracy on all previous tasks is recomputed after every task on a heldout test set and averaged; error bars denote 95% confidence intervals. The sticky gradient method is present in both of the two most accurate graphs. The control simply refers to a network initialized with a standard Kaiming initialization and no L1 regularization (He et al., 2015).

### 3.4. Shared Information

Sets of real world image-processing tasks almost never have uncorrelated inputs. As such, we evaluate the method on variations of the PMNIST dataset for which a proportion of pixels are not permuted. We compare a baseline 2000 ReLU ANN with Kaiming initialization against a network of the same architecture equipped with the *sticky gradient* method for different levels of shared information (He et al., 2015). Batch size was set to 50 and each network was trained for a single epoch on every task and optimized via Adam.

We find that the *sticky gradient* method does not lose a benefit until the proportion of shared information reaches 0.6 (Figure. 4). This illustrates the effectiveness of our method on correlated tasks akin to real-world scenarios in which agents may be asked to complete tasks which leverage knowledge gained from previous tasks. However, it also points out a trade-off that our method makes between expressivity and retention. As the *sticky gradient* network trains, more weights get caught in the dip of the weight-plasticity function and are unable to shift in response to new examples that may improve performance.

**Figure 4:**
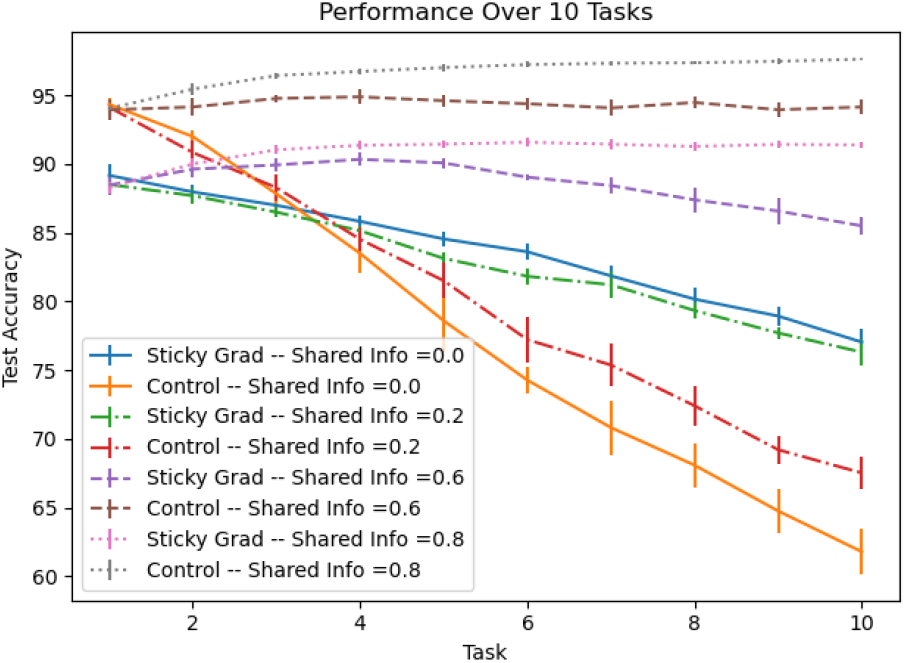
Accuracy profiles from runs for 10 random seeds for different levels of shared information; error bars denote 95% confidence intervals. Our method retains a benefit over the control until the amount of shared information reaches 60%. After this point, the *sticky gradient’s* inability to modify frozen weights for similar tasks outweighs its ability to retain previous knowledge.

### 3.5. Partial Sticky Gradient

In the *sticky gradient* algorithm, the base learning rate is zero. Intuitively, this means that synapses near the attractors are rendered completely immobile. We test the assumption that this configuration is optimal by using a slightly modified version of *Eq. 1* with two parameters in place of *λ*. In this way, we allow the meta-learner the freedom to choose parameters for which the minimum learning rate is not 0.

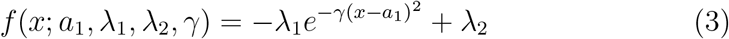

We meta-search over the adapted set of hyperparameters *τ* = {*a*_1_, *µ, σ, γ, ζ, λ*_1_, *λ*_2_} for 200 trials to produce the partial *sticky gradient* parameters given by *Table 2*. The meta-learned values of *λ*_1_ and *λ*_2_ are significantly different, creating a minimum learning rate multiplier of 0.43. The rest of the resulting parameters are strikingly similar to those of the first meta-search (*Table 1*), significantly differing only in the magnitude of *γ*.

**Table 2:** Learned hyperparameters from 200 trial meta-search. Interestingly, the metasearch yields markedly different values of *λ*_1_ and *λ*_2_, underlining the potential importance of a non-zero learning rate near the attractors.

| $a_1$ | $\mu$ | $\sigma$ | $\gamma$ | $\zeta$ | $\lambda_1$ | $\lambda_2$ |
| --- | --- | --- | --- | --- | --- | --- |
| 0.252 | 0.00320 | 0.0191 | 2.18 | 0.000386 | 4.51 | 4.94 |

We compare the efficacy of the *partial sticky gradient* method with hyperparameters *τ* with the control and sticky gradient using one-hidden-layer 2000 ReLU networks trained for a single epoch for each task of a 10 task sequence and optimized using Adam with a base learning rate of 1e-4. The *partial sticky gradient* method performs on par with EWC at 10 tasks in terms of both average and previous task performance (*Figure 5a/5b*). This performance implies that partially frozen weights being allowed some freedom to move in response to new information can be beneficial, provided that it keeps them sufficiently close to the attractor well, however this benefit is small.

**Figure 5:**
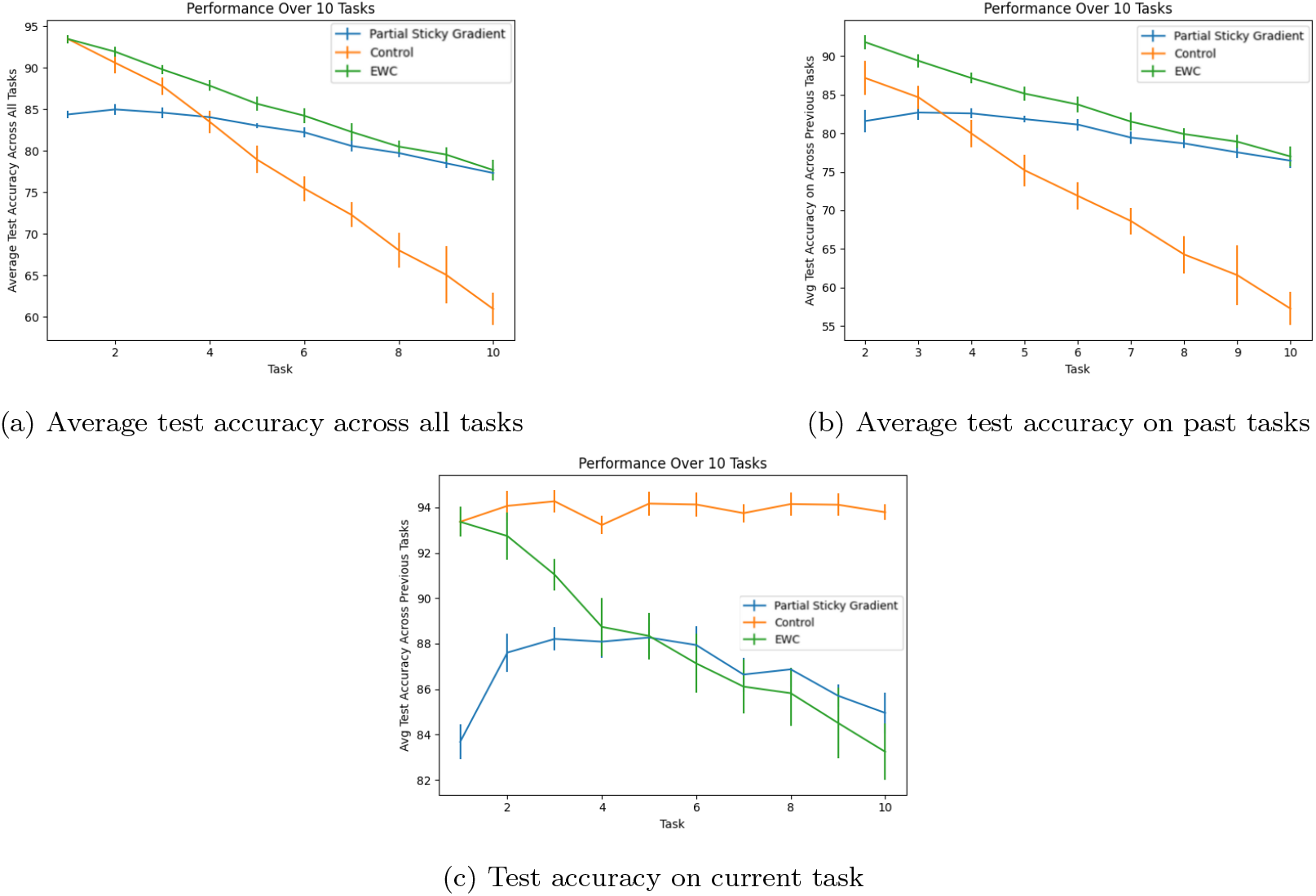
(a) Comparison study detailing the test accuracy of trials under 10 different random seeds. Accuracy on all tasks is recomputed after every task on a held-out test set and averaged; error bars denote 95% confidence intervals. At 10 tasks, EWC and the *sticky gradient* method outperform the control by an average margin of 19%. The partial *sticky gradient* method and EWC perform almost equally well at 10 tasks. (b) Average test accuracy on all tasks, not including the current one. Accuracy is computed on all previous tasks after every task except the first and averaged; error bars are 95% confidence intervals. Again, our method is on par with EWC in this setting at 10 tasks and both methods outperform the control by an average of 23%. (c) Test accuracy on current task; error bars denote a 95% confidence interval. EWC has a small decrease in the ability to learn new tasks over the 10 task process. The partial *sticky gradient* method retains a similar receptivity to new tasks over this horizon, but starting from a lower initial task learning. Both continual learning methods lag the control by about 9% on this metric after 10 tasks.

### 3.6. Weight Dynamics

It is also instructive to look at the dynamics of the weights during training. We expect both variations of the *sticky gradient* weights to behave quite differently than those of the control over the course of training. We discover that this is indeed the case – the eventual weights for the full *sticky gradient* method are essentially trimodal. There is a group of weights near zero, another near the dip of the weight-plasticity function, and yet another contingent of inhibitory weights. Broadly, the partial *sticky gradient* method seems to be quite similarly distributed. The partial distribution is composed of an inhibitory group that is less obvious due to the decreased distance from the zero-peak, a zero-peak, and a less-concentrated mass of weights near the attractor value. In contrast, both the control and EWC exhibit clear unimodality around 0 (*Figure 6*).

**Figure 6:**
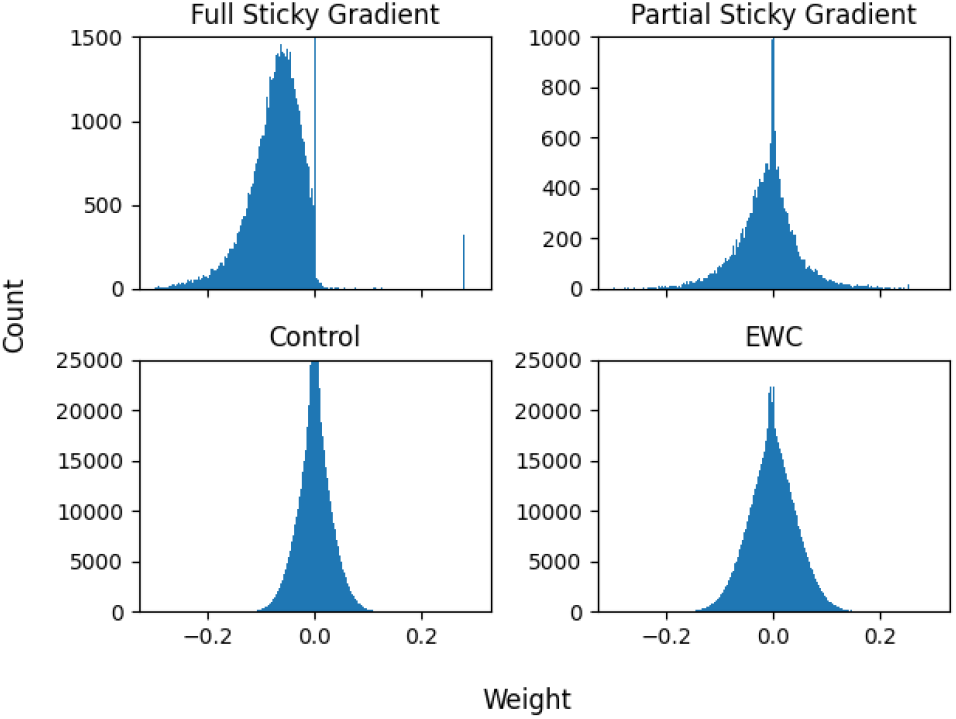
Weight distributions from Kaiming initialized control, EWC, and full/partial *sticky gradient* networks. We truncate the y-axis for the sticky gradient histograms to ensure that the non-zero peaks remain visible. The EWC and control networks seem to exhibit similar patterns of unimodality around 0, differing only in width. The full *sticky gradient* network produces an interesting distribution of weights with local modes near -0.06, 0, and 0.279 (the attractor value). The partial *sticky gradient* network is also multimodal with local maxima near 0 and 0.252 (the attractor value). As expected, weights are far less concentrated near the attractor in the partial model.

In essence, both the full and partial *sticky gradient* methods seems to encourage the freezing of synapses near the attractor, and the masked gradient multiplier induces a sort of symmetry by increasing the rate at which faraway weights move away from the attractor. This interplay allows for a balancing in which a contingent of inhibitory neurons arises to combat the excitatory effect of their positive counterparts near the attractor. In support of this, we note that the *partial sticky gradient*, which has fewer positive weights, also has fewer strongly inhibitory weights – despite the similarity between in the magnitudes of *λ, λ*_1_, and *λ*_2_ (*Figure 6*). We hypothesize that concentration of information in these relatively few parameters enables the observed superior retention of knowledge from previous tasks.

## 4. Discussion

We show that groups of synapses endowed with even the simplest prior measures of weight-dependent plasticity can be used to enable continual learning in ANNs. Specifically, we show that it is possible to meta-learn parameters for a flipped gaussian function mapping weights to plasticity that aids in continual learning.

Our method bears similarity to Zenke et al.’s SI and Kirkpatrick et al.’s EWC in that we aim to regulate plasticity in response to some measure of weight importance. However, both of these methods do this explicitly by keeping track of this measure of weight importance and penalizing it in the loss (Kirkpatrick et al., 2017; Zenke et al., 2017). In contrast, we allow the meta-learner to implicitly come up with a region of a high weight importance based on training dynamics by meta-learning parameters for the weight-plasticity function.

Of all the plasticity-related methods for continual learning, ours is potentially the easiest to justify biologically. While SI requires complicated explanations about the plausibility of a separate store of importance in synapses, we do not make use of extra synaptic dimensions. (Montgomery and Madison, 2002). Indeed, we show that it is possible to effectively alleviate catastrophic forgetting simply by creating a global mapping of synapse strength to importance. This mapping is eminently biologically plausible – there is already evidence that plasticity is activity dependent (Cingolani et al., 2008) and likely plasticity is also weight dependent and therefore that our method is biologically grounded.

The weight distributions created by the *sticky gradient* methods are actually reminiscent of recent work concerning distributions of synaptic weights in mouse connectomes. In this study, the authors find that, controlling for cell type, synapse strength can be described as a bimodal mixture of lognormal distributions (Dorkenwald et al., 2019). Indeed, the weight distributions produced by the *sticky gradient* method show marked multi-modality due to pooling near the attractor. While this does not exactly match the bimodality described in the study, the similarity that exists even under these radically different conditions undeniably adds to the biological plausibility of our method (Dorkenwald et al., 2019).

Our work points to the effectiveness of this particular brand of biological prior on enabling continual learning in artificial neural networks. More broadly, our research highlights the ability of artificial neural networks to benefit from even incredibly simple plasticity-related priors. This opens a number of avenues for further work in which a broader class of system-wide plasticity-related priors can be leveraged to effectively manipulate training dynamics for a wide variety of learning environments.

## 5. Acknowledgments

This work has been performed as a part of the Allen Institute summer internship project. We would like to thank the entire Mindscope Modeling & Theory group at the Allen Institute for their insight and suggestions. We thank the Allen Institute founder, Paul G. Allen, for his vision, encouragement, and support.

## 6. Appendix

### 6.1. Permuted MNIST Experiments

We use fully-connected feedforward 2000 ReLU multi-layer perceptrons for classification on every task. To enable the control networks to better retain knowledge from previous tasks, we train each model for a single epoch on each dataset. A full list of fixed hyperparameters is given by *Table 2*. All networks trained with the *sticky gradient* method meta-optimize parameters on a validation set, disparate from the training set. All networks are tested on held-out test sets over 10 random seeds. Additionally, we use an Adam optimizer every epoch but reset its state after every task to prevent differing initial learning rates per task.

### 6.2. Meta-Learning

We use a simple TPE-mediated search to find *ϕ* and *τ* under the following conditions over 30 and 200 trials, respectively, using the search ranges in *Table 3* (Bergstra et al., 2*011*). Parameters were assumed to have a uniform prior over the log domain of the specified ranges. All networks are trained using hyperparameters *ϕ* (full) or *τ* (partial) for a single epoch per task for 10 tasks using the Adam optimizer with learning rate 1*e* − 4.

**Table 3:** Hyperparameters for PMNIST Experiment

|  |  |
| --- | --- |
| Base Learning Rate | $10^{-4}$ |
| Epochs per Task | 1 |
| Hidden Layer Width | 2000 |
| Depth | 1 |
| N. Hidden Layers | 1 |
| Optimizer | Adam |

**Table 4:** Ranges for meta-search

| $a_1$ | $\mu$ | $\sigma$ | $\gamma$ | $\zeta$ | $\lambda$ |
| --- | --- | --- | --- | --- | --- |
| 0.01 - 10.0 | 1e-3 - 8.0 | 0.01 - 7.0 | 0.01 - 7.0 | 1e-7 - 1e-3 | 4.0 - 10.0 |

